## Supplementary Materials for "Chromatin Regulates Bipartite-Classified Small RNA Expression to Maintain Epigenome Homeostasis in Arabidopsis"

**Additional File 4:** Supplementary Figures

**Table of contents**

**Figure S1.** Characteristics of embryonic 24-nt siRNAs and their similarities across samples

**Figure S2.** siRNA dynamics and characteristics

**Figure S3.** Benchmarking low-input methylC-seq, methylomes and cell-cycle transcripts

**Figure S4.** Size-based partitioning of heterochromatic TEs and small RNA-directed methylation

**Figure S5.** TE-derived siRNA accumulation and association with chromatin

**Figure S6.** TE-derived siRNAs in methylation mutants

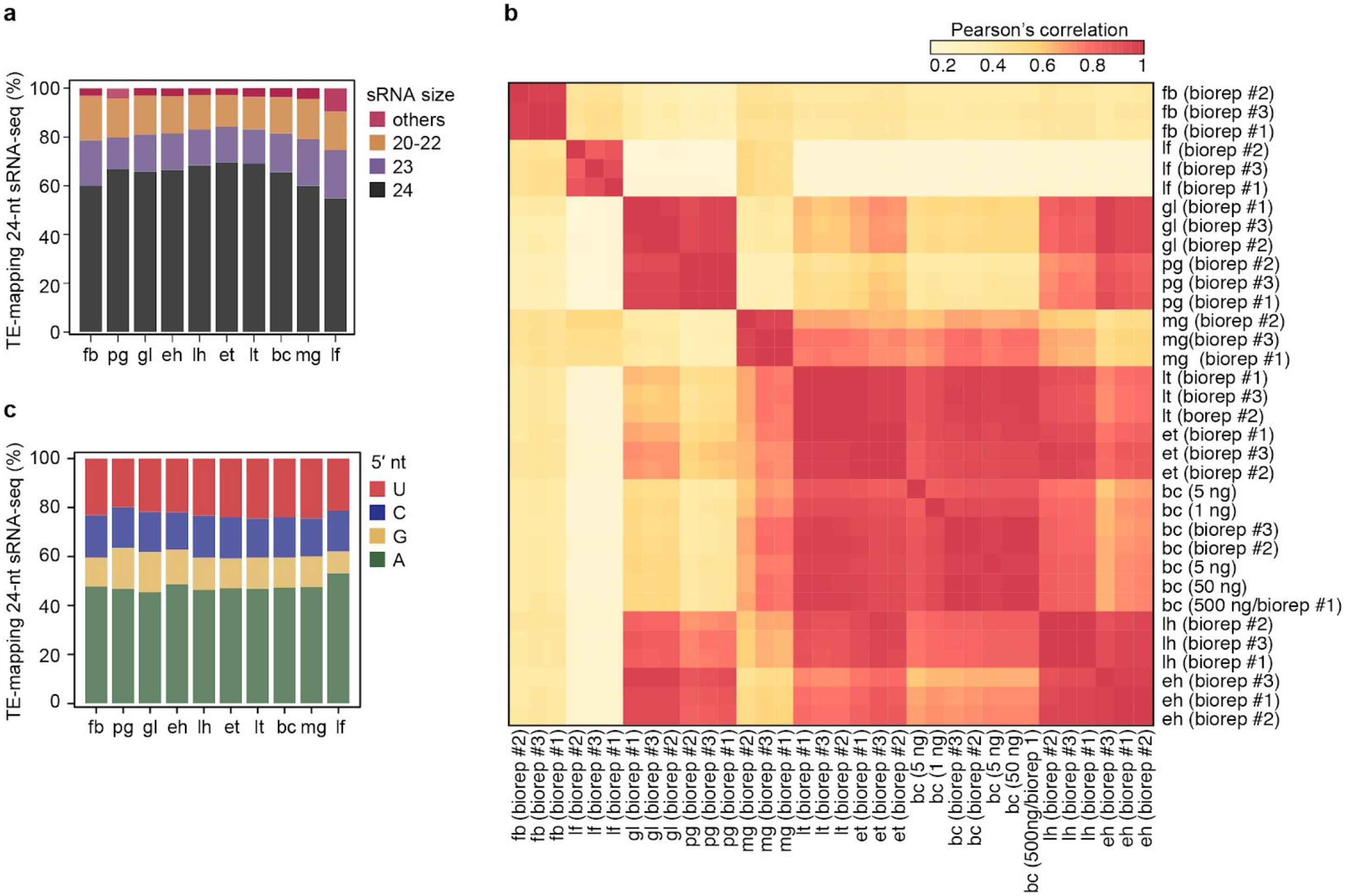


**Fig S1.** Characteristics of embryonic 24-nt siRNAs and their similarities across samples, related to Fig 1. **a** Stacked bar chart depicting the percentage (%) of reads mapping to TEs according to their sizes. Others include 18, 19 and 25-30 bases. **b** Heatmap of Pearson's correlation coefficients of 24-nt siRNAs among different biological replicates and tissue types. **c** Stacked bar chart showing the percentage (%) of 24-nt siRNAs with various 5′-most nucleotides in floral bud, embryonic and leaf tissues. Nucleotides are color-coded according to the key. Sample labels are as shown in Fig 1a.

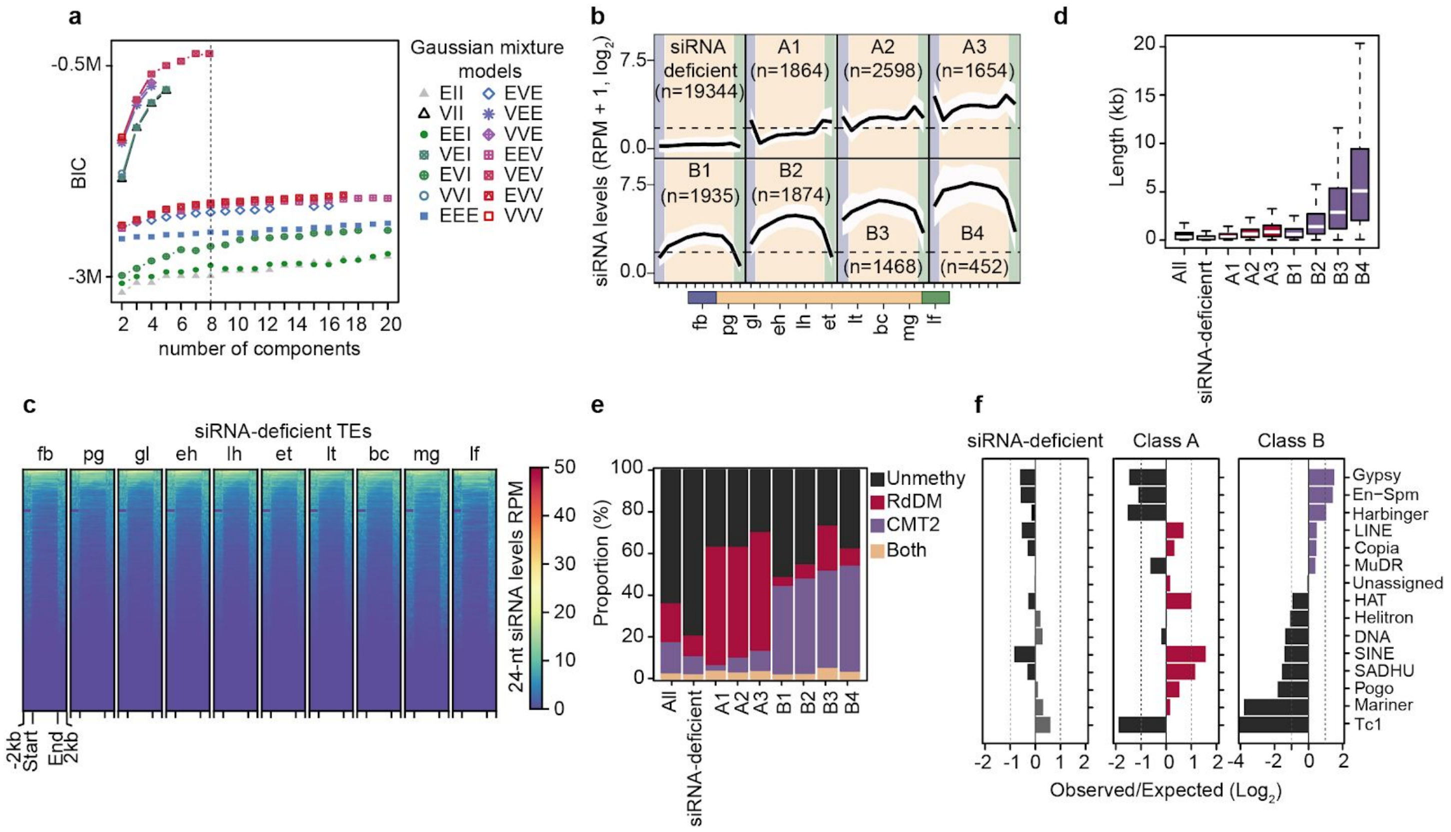


**Fig S2.** siRNA dynamics and characteristics, related to Fig 2. **a** Bayesian information criterion (BIC) of various Gaussian mixture models of TE-derived 24-nt siRNAs as determined by *mClust*. EII, Equal volume, Equal shape, VII, Variable volume, Equal shape; EEI, Equal shape, Equal volume, coordinate axes orientation; VEI, Variable volume, Equal shape, coordinate axes orientation; EVI, Equal volume, Variable shape, coordinate axes orientation; VVI, Variable volume, Variable shape, coordinate axes orientation; EEE, Equal volume, Equal shape, Equal orientation; EVE, Equal volume, Variable shape, Equal orientation; VEE, Variable volume, Equal shape, Equal orientation; VVE, Variable volume, Variable shape, Equal orientation; EEV, Equal volume, Equal shape, Variable orientation; VEV, Variable volume, Equal shape, Variable orientation; EVV, Equal volume, Variable shape, Variable orientation; VVV, Variable volume, Variable shape, Variable orientation. **b** Line graphs illustrating 24-nt siRNA levels from siRNA-deficient TEs, as well as originally defined three class A and four class B TEs in embryonic and post-embryonic tissues. Dashed lines represent the detection criteria used to select TEs yielding siRNAs (2 RPM, reads per million genome-matching reads). The number of TEs belonging to each class are indicated. Polygons represent the standard deviation of mean 24-nt siRNA levels. fb, floral buds; pg, preglobular; gl, globular; eh, early heart; lh, late heart; et, early torpedo; lt, late torpedo; bc, bent cotyledon; mg, mature green; lf, leaf. **c** Heat map showing 24-nt siRNA RPM from siRNA-deficient TEs across development. siRNA levels from ±2-kb of TEs are color-coded according to the key, and samples are labelled as in Figure 1A. Rows were ordered based on total 24-nt siRNA levels. **d** Boxplot of TE lengths for *mClust*-defined groups including all annotated, siRNA-deficient, class A and class B TEs. Thick horizontal bars indicate medians, and the top and bottom edges of the box indicate the 75th and 25th percentiles, respectively. kb, kilobases. **e** Stacked bar charts illustrating proportion of TE subclasses that are CHH hypomethylated in *drm1*/*drm2* (RdDM; red), *cmt2* (CMT2; purple), both *drm1*/*drm2* and *cmt2* (both; yellow) or which were not methylated (unmethylated; black) in leaves. **f** Enrichment of TE families observed in siRNA-deficient, class A or class B groups relative to their respective genomic backgrounds (observed/expected; $log_2$).

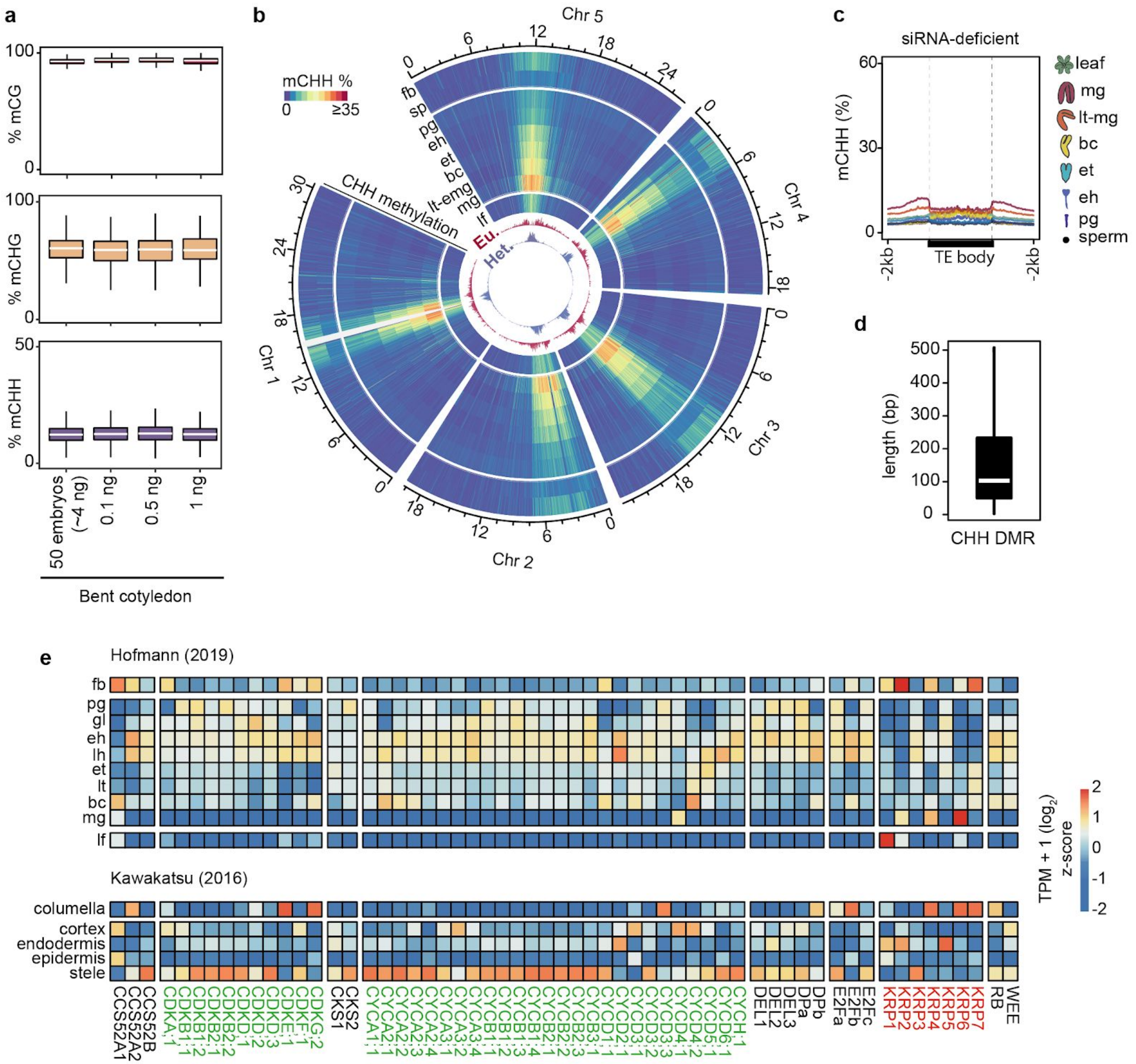


**Fig S3.** Benchmarking low-input methylC-seq, methylomes and cell-cycle transcripts, related to Fig 3. **a** Box plots of average weighted CHH methylation percentages of TEs with at least 5 informative cytosines ≥3× coverage in either CG (*top*), CHG (*middle*) or CHH (*bottom*) contexts from various amounts of input genomic DNA. Labels on x-axis indicate whether these were from 50 bent cotyledon embryos (~4 ng), or a dilution series of genomic DNA from bent cotyledon embryos. Thick horizontal bars indicate medians, and the top and bottom edges of the box indicate the 75th and 25th percentiles, respectively. **b** Circos heat map representing average weighted CHH methylation of euchromatic (Eu.; red) or heterochromatic (Het.; purple) TEs in reproductive, embryonic and vegetative tissues across the five Arabidopsis chromosomes (Chr). Unmapped region in chromosome 1 (Chr 1) is shaded in white. fb, floral bud; sp, sperm; pg, preglobular; eh, early heart;

lh, late heart; et, early torpedo; bc, bent cotyledon; lt-mg, late torpedo-to-early mature green; mg, mature green. **c** Metaplot of average weighted CHH methylation percentages of siRNA-deficient TEs sperm, embryos and leaves. Abbreviations are as in (B). **d** Box plot of CHH developmental DMR lengths, and as described in (A). **e** Heat maps of relative transcript levels from cell-cycle related genes across floral buds, embryos and leaves (*top*) or individual root cell types (*bottom*). Cell-cycle promoting and inhibitor gene labels are color-coded in *green* and *red*, respectively. Relative transcript levels are shown as z-scored TPMs according to the key.

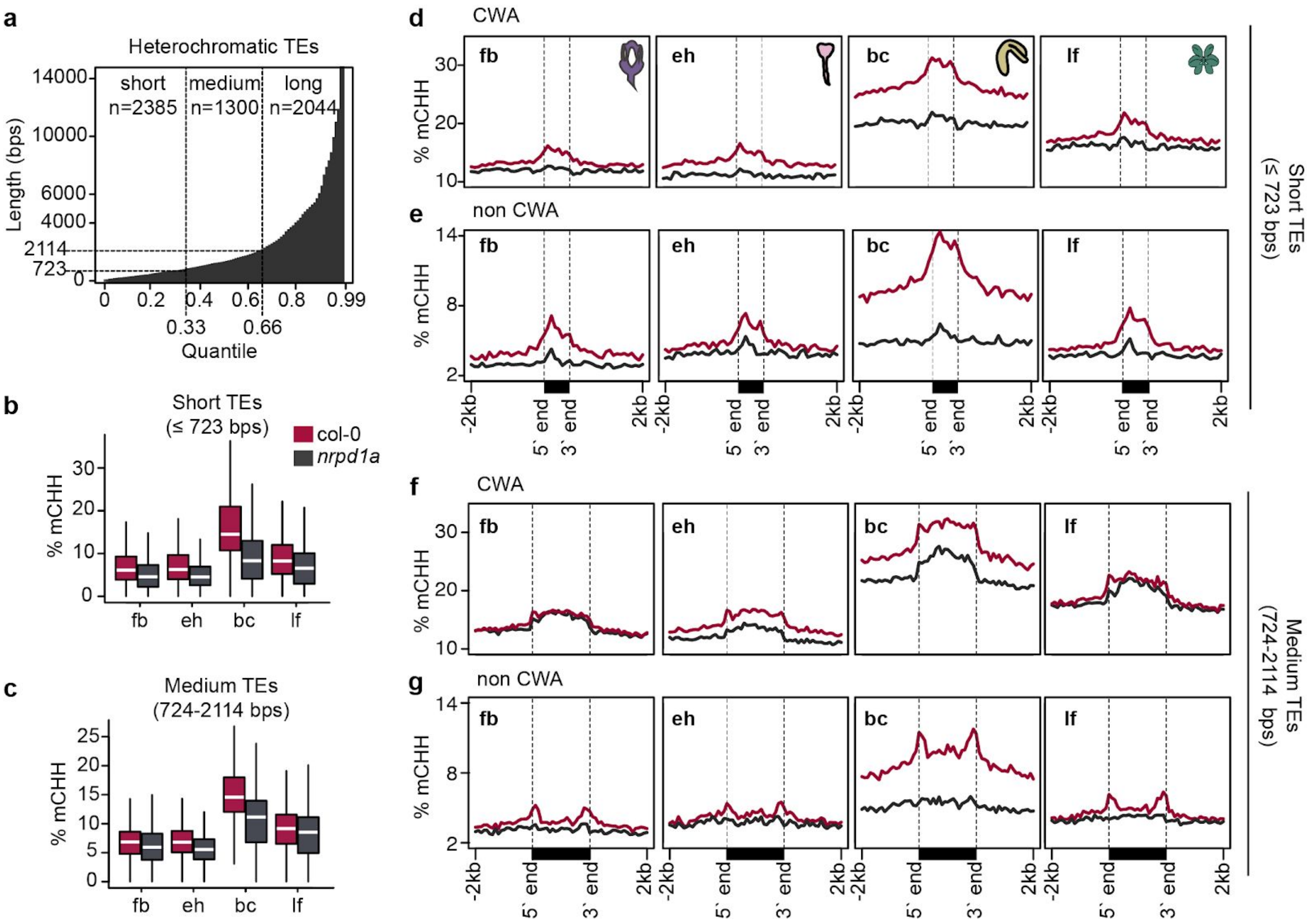


**Fig S4.** Size-based partitioning of heterochromatic TEs and small RNA-directed methylation, related to Fig 4. **a** Histogram of heterochromatic TE lengths across quantiles, which were used to separate TEs into short, medium or long classes. Vertical and horizontal dashed lines represent quantiles and sizes used for classifications. **b** and **c** Boxplots of CHH methylation levels in wild type and *nrpd1a* mutant floral buds (fb), early heart embryos (eh), bent cotyledon embryos (bc) and leaves (lf) for short (b) and medium (c) heterochromatic TEs. **d-f** Metaplots of average weighted CHH methylation percentages of short (d and e) and medium (f and g) heterochromatic TEs in either CWA (d and f) or non-CWA (e and g) contexts.

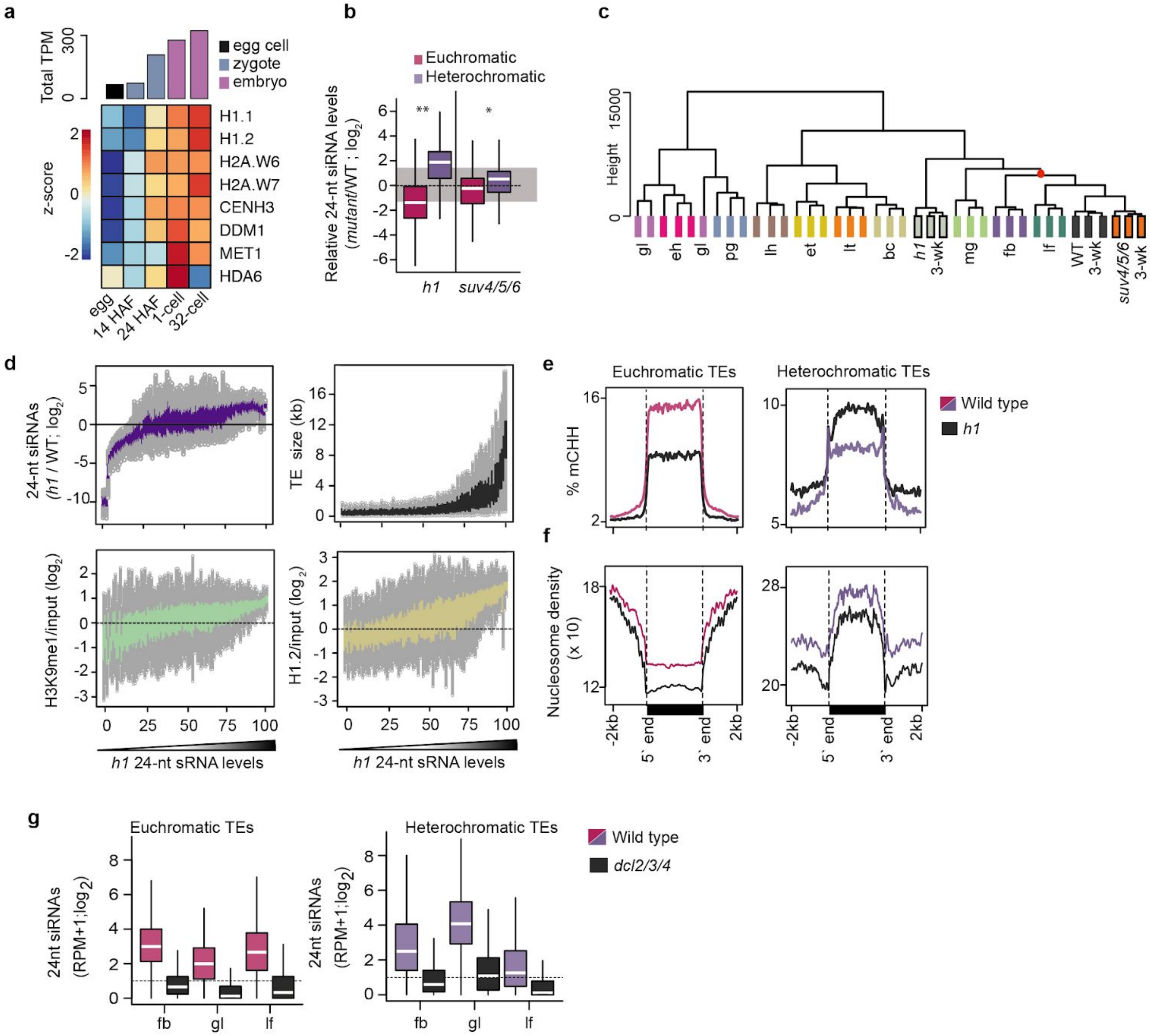


**Fig S5.** TE-derived siRNA accumulation and association with chromatin, related to Fig 5. **a** Bar chart depicting total (*top*) and heat map of individual (*bottom*) transcript levels from genes involved in heterochromatin formation in eggs, zygotes 14 or 24 hours after fertilization (HAF), and embryos at 1-cell and ~32-cell stages. **b** Boxplots of relative 24-nt siRNA levels overlapping euchromatic (red) or heterochromatic (purple) TEs in *h1* or *suv4*/*5*/*6* mutants relative to wild-type leaves. *P* values < 0.001 and <0.0001 based on Mann-Whitney U test 24-nt siRNA differences between wild-type and mutant tissues are represented by * and **, respectively. **c** Dendrogram based on hierarchical clustering of Euclidean distances of TE-derived 24-nt siRNAs from embryonic and post-embryonic tissues (key). Breakpoint of embryonic and non embryonic tissues are indicated by a red dot. **d** TEs were divided into percentiles, ordered based on their 24-nt siRNA levels in *h1* mutants (as in Fig 5E-H), and plotted according to 24-nt siRNA levels in *h1* relative to wild type, TE length, enrichment of H1.2 or H3K9me2. **e** and **f** Metaplots of average weighted CHH methylation percentages (E) and nucleosome occupancy (F) of euchromatic (*left*) and heterochromatic (*right*) TEs.**g** Boxplots of 24-nt siRNA levels of euchromatic (*top*) and heterochromatic (*bottom*) TEs in wild type and *dcl2*/*3*/*4* floral buds (fb), globular embryos (gl) and leaves (lf) (key).

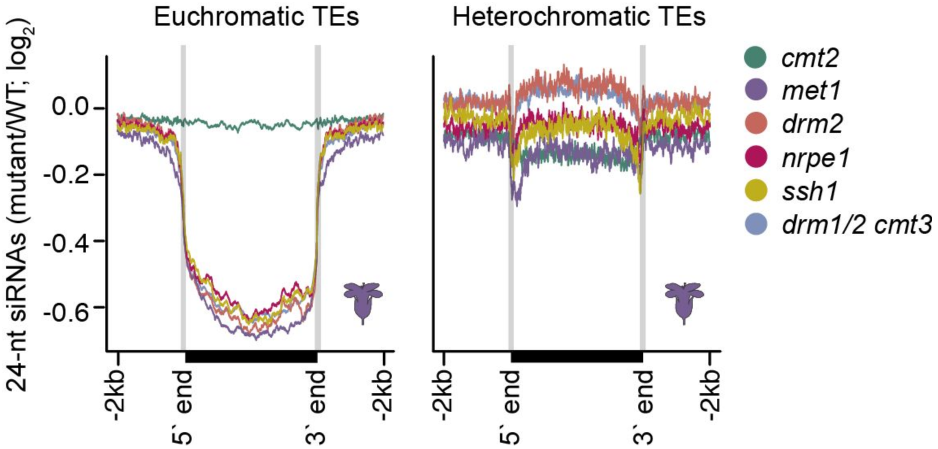


**Fig S6.** TE-derived siRNAs in methylation mutants, related to Figure 6. Metaplots of 24-nt siRNA levels overlapping euchromatic (*left*) and heterochromatic (*right*) TEs in various RdDM and related mutants. Annotated 5′ and 3′ ends are labelled at the bottom.
